# Pollinator Plant Network Interactions of Bees (Hymenoptera: Anthophila) in a California Urban Garden

**DOI:** 10.64898/2026.05.13.724999

**Authors:** Isabel Navarro, Nina A. Sokolov

## Abstract

Amongst ever increasing habitat loss and compounding environmental stressors, urban pollinator gardens can provide refugia and support native bees. However, gardens are rarely assessed for their effectiveness in providing adequate habitat. This study focuses on a recently established local campus pollinator garden in California. Using diversity observations from the early 2000s garden and non-lethal sampling techniques, we characterized visitation frequencies between flowers and urban bees. We generated a bipartite interaction network and found that despite the garden’s relatively small size, the network showed greater specialization and modularity, as well as lower nestedness than expected by chance. Across 12 flower species, we observed two non-native pollinators, the western honey bee (*Apis mellifera*) and the alfalfa leafcutter (*Megachile rotundata*), along with at least nine native bee species across three families (Apidae, Halictidae, Megachilidae). Although this garden was designed for native bees, honey bees represented over 80% of visitation events and thus suggests that the garden needs adjustments to provide more forage for native bees. Honey bees occupied the most central and connected position in the network, followed by native sweat bees (Family: Halictidae) which both had generalist patterns. Whereas other bees showed evidence of specialization (Genus: *Mellisodes*). We found a significant positive relationship between inflorescence count and honey bee abundance but not with native bee abundance. To better steward native bees that possess a range of floral specializations, managers may consider increasing plant species diversity rather than relying on a few but highly abundant flowering plants.

## Introduction

Insect pollinators play a vital role in maintaining wildflower diversity and supporting food systems. Pollinators provide key ecosystem services that support animal and plant populations, and it is estimated that 85% of flowering plants rely on bee pollination (Hernandez et al. 2009a). The most commonly managed species used for food crops is the western honey bee (*Apis mellifera* (Linnaeus, 1758), which pollinates a third of the food in the global agricultural landscape (Daniels et al. 2020). In other words, the current agricultural system in the United States is heavily reliant on the pollination services from managed, non-native *A. mellifera* for crop production. Anthropogenic activities such as urbanization have significantly degraded natural habitats for wild and native bees. Conversion of wildland habitat for human use reduces foraging resources and nest-site availability for bee populations (Hernandez et al. 2009). Agrochemicals, such as pesticides and insecticides, that are applied to farmlands harm bee populations, diminishing reproduction, immunity, memory, and navigation (Chmiel et al. 2020). Managed honey bee *A. mellifera* hives have been shown to create ecological pressures on native bee species by acting as competition for floral resources (Page & Williams 2023) and increasing the potential for disease transmission (Graystock et al., 2016).

Despite environmental threats to bee populations, some native bee species have adapted to anthropogenic changes and can thrive in residential areas, referred to as ‘urban bees’ (Hernandez et al. 2009). A large-scale study in the UK found that residential/community gardens were pollinator hotspots (Baldock et al. 2019). Studies conducted in several major cities, including Berkeley, California, show that urban gardens can support diverse bee populations (Philpott et al., 2025; Hall et al. 2017). Urban gardens can attract and support bee species if there is available nesting habitat and if floral resources are diverse in phenology, with blooms spanning the entire bee pollination season (Wojcik et al. 2008). To successfully support bee populations, urban gardens must include appropriate plant species in sufficient quantities (Philpott et al., 2025; Baldock et al., 2019; Frankie et al. 2009,).

Gardens can act as a ‘refuge’ for insect pollinators, aiming to recreate a portion of the resources found in wildland habitats (Hall et al. 2017). for In fact, bumblebees (genus *Bombus* Latrielle, 1802) may exhibit higher species richness in gardens than in natural habitats (McFrederick & LeBuhn 2006; Hernandez et al. 2009). Studies across ten California cities found that native bees visit both native and non-native plants, but non-native bees visit non-native plants more often. The use of non-native and native plants with differing flowering phenology can support bees throughout the full pollinator season, such as the social bee yellow-faced bumblebee *(Bombus vosnesenskii* (Radoszkowski, 1862)) (Frankie et al. 2009). Urban gardens can support wild bee biodiversity; California urban garden sampling for a period of over 15 years has documented approximately 400 bee species, representing 20% of California’s 1,600+ bee species (Frankie et al. 2019).

The University of California (UC) Urban Bee Lab, located at the Oxford Tract campus research farm, maintains an experimental urban garden of California native and non-native plants to attract and support native wild bee biodiversity. Historically, this was maintained as a garden plot from 2003-2009, with approximately 85% California native plants, to assess emerging patterns of diversity and seasonality (Wojcik et al. 2008). This garden was approximately 180 m^2^ and established current understandings of urban bees known to be found in Berkeley. In 2004, there were 78 plant species in the garden, aggregated into 1.5 m^2^ patches. The plot plants were selected to provide continuous, consistent blooms during the known phenology of the local bees (Wojcik et al. 2008). A 2004 study reported that 32 bee species, 17 genera, and five families utilized the garden for floral resources. A 2009 study found 82 urban bee species in Berkeley, including the naturalized but non-native honey bee (*A. mellifera)* and the alfalfa leafcutter bee (*Megachile rotundata* (Fabricius, 1793)) (Frankie et al. 2009). In the late 2010s, that garden was decommissioned, and a new urban bee garden at the research farm was established. Similarly to the original garden, the new plant selection provides pollen, nectar, and habitat resources for ground-nesting and wood-nesting bees. While studies were conducted in Berkeley from 2000 to 2010, there are currently no studies of the new pollinator garden. By understanding how wild bees use native and non-native plant resources provided by urban spaces, habitat gardening practices can be refined to best support native bee biodiversity. Plant pollinator networks offer us a key tool in understanding the complex interactions between the flowers we choose to cultivate in gardens and their bee pollinators.

In this study, we surveyed bee diversity in an urban garden on the University of California, Berkeley campus at the Oxford Tract research facility. We wanted to understand the plant-pollinator interactions and community dynamics of the species in the garden over time, including how much *A. mellifera* dominated the interactions, and how the garden’s bee diversity compared to past records. To do this, we created a bipartite plant-pollinator network to describe interactions between flowers and native bee species in this urban garden. Bipartite plant-pollinator networks visualize pollination interactions between plants and bees as interconnected nodes, weighted by species abundance and the number of observations (Maurer et al. 2024). Interaction networks can be used to understand the structure of community interactions among plants and pollinators, as well as between different pollinator species. This enables a deeper understanding of the mechanisms underlying an urban garden environment and informs future stewardship strategies for floral plantings (De Sousa Perugini et al. 2025). Network metrics give insight into the vulnerability of urban communities to disturbances, specialization of interactions, and the roles of particular species. Plant-pollinator networks for other urban garden spaces have identified plant morphological characteristics and floral abundance as key drivers of interactions (De Sousa Perugini et al. 2025). This network will enable us to assess the garden’s effectiveness through data visualization and improve the native pollinator garden design through ecologically informed stewardship applications.

(1) We hypothesize that the urban garden displays a non-random interaction network with high modularity corresponding to some groups of bees interacting with one another more than others. Next, we predict that native bees would show higher levels of floral specialization while non-native bees would demonstrate generalist foraging patterns. (3) Because the urban garden has been curated to best support native bees, we predict that less than 50% of bee visitors would be non-native species like *A. mellifera*. (4) Finally, we hypothesize that there will be an overall loss in native bee biodiversity at the Oxford Tract field site since the 2004 study due to a reduction in the flower selection and diversity in the garden.

## Methods and Materials

### Field Site

The garden measures approximately 108.96 m^2^ and is located in the parking lot median of the Oxford Tract Research Facilities at the University of California, Berkeley campus in Berkeley, California, U.S.A [37.875676, -122.267580]. Neither mulch nor grass is used in order to leave the soil bare. Instead, naturally occurring leaf litter is used to support ground-nesting bees. Decaying trunks provide habitat for wood nesting bees such as carpenter bees (genus: *Xylocopa* Latreille, 1802) and small carpenter bees (genus: *Ceratina* Latreille, 1802). Plants were introduced into the urban garden space, whether native or non-native, typically in groups of three and were pruned and watered as needed. All plants were identified upon purchase from the nursery gardens.

### Data Collection

Plant–pollinator observations were conducted in a single urban community garden throughout the month of July 2025. Non-lethal collection methods were used to minimize impact and conserve the garden’s limited native bee population. As this study aims to report interaction dynamics of the entire community, lethal collection could disrupt accurate measurements on subsequent survey days. Our methods follow “compassionate collecting”, avoiding unnecessary disturbance to species in their natural habitats for conservation purposes (Byrne, 2023). We found non-lethal samples to be sufficient in identifying bee species to genus or species on account of the existing natural history records of the Oxford Tract’s urban bees in published literature (Frankie et al. 2009) and specimen collections. Photography as a collection method increases access to specimens by allowing them to be viewed online, thereby removing the need for visitors to private collections.

Only actively blooming plants in the garden space were included in the study. All flowering plant species in the garden were monitored, and all of the bees contacting floral reproductive structures were recorded to the lowest possible taxonomic level (Fig. 1). Over three survey days, bees were observed and collected from 12 blooming plant species at the urban garden between 10:00 and 16:00. As different bee species forage at various times of day (Frankie et al. 2009), surveys were divided into 2-hour periods, with three time points: (1) morning, (2) midday, and afternoon, to capture the range of foraging phenology. Each plant species was observed daily to generate a complete profile of bee pollinator interactions across the three time points. In total, sampling lasted across three days. Bee visitors were observed on each plant for 10 minutes, and the number of pollination events per bee genus was recorded as a measure of visitor abundance. Following abundance measurements, representative bee species were collected using aerial nets for 15 minutes, and each individual was placed into a labeled centrifuge tube. Tubes were then put into an ice chest for 15-30 minutes for temporary sedation. The sedated bees were photographed for identification purposes and then released in a shady area within 100 meters of the garden. Abundance and collection methods were repeated across four different plants over time. Bee traits data, including socialness, mean proboscis length, foraging habits, native range, and phenology, were collected from the literature. During each survey day, the number of blooming flowers on a plant was recorded as an abundance measurement. The total number of blooms across all individuals of each plant species was counted on each field day. Bloom counts for each species across all field days were averaged to obtain the mean inflorescence count. Corolla length and depth measurements for 10 individual blooms from multiple plants were recorded with callipers and averaged (Pélabon et al. 2013). Collection days were scheduled as close as possible, while accounting for windy, overcast, and rainy conditions.

**Figure 1.**
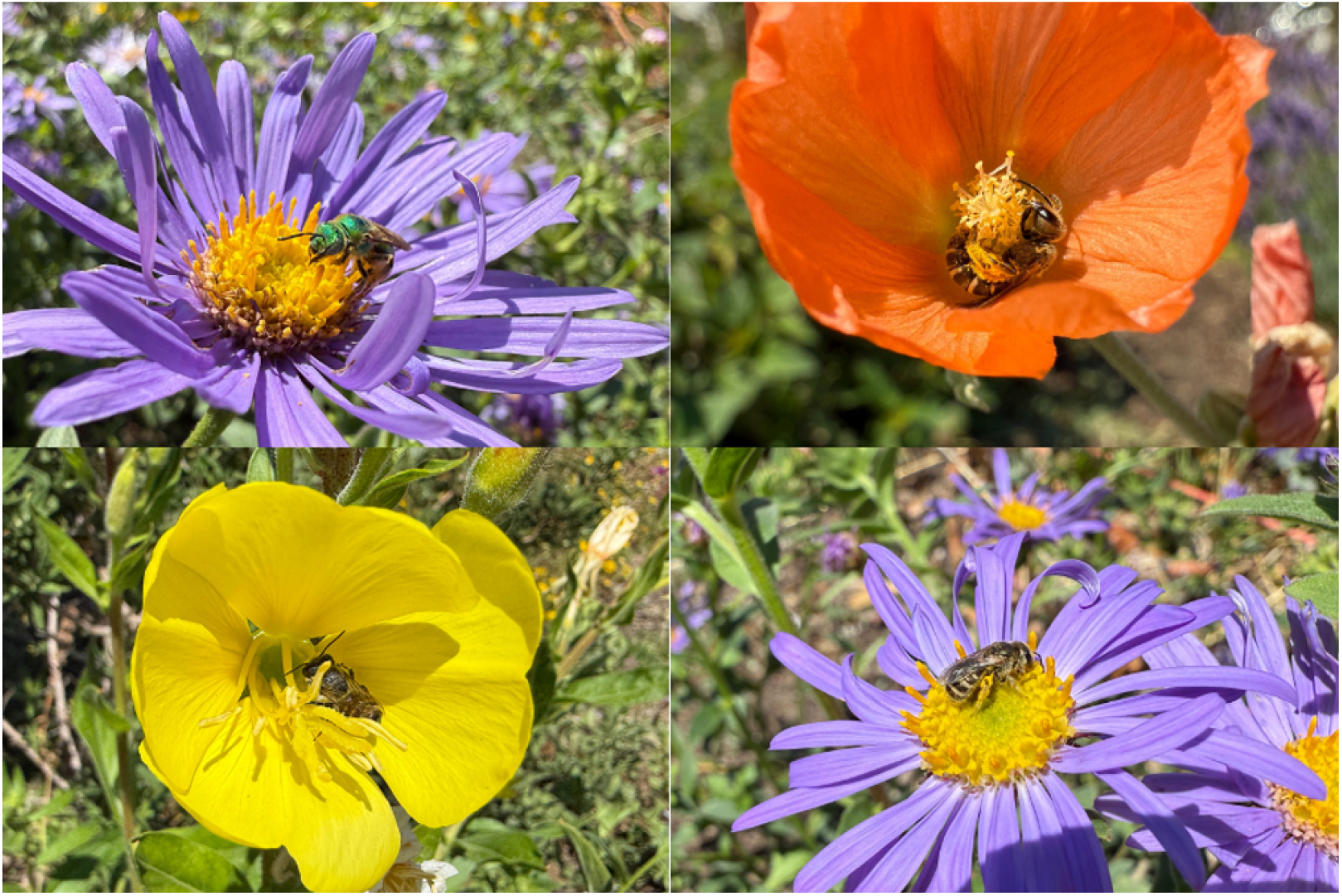
Photographs of bee visitors on blooming flowers in the Oxford Tract pollinator garden field site across multiple observation days. *Agapostemon subtilior* on *Aster x frikartii* (top left), Halictidae on *Sphaeralcea ambigua* (top right), *Megachile* sp. on *Oenothera* sp. (bottom left), Halictidae on *Aster x frikartii* (bottom right).

### Bee Identification

Bees were identified in the field during abundance measurements to the highest taxonomic level possible (genus or family). Photographed bee representatives were identified to genus by comparisons to previously identified species from Gordon W. Frankie (GWF) collections from Berkeley, 2001-2015 and validated using published taxonomic dichotomous keys. The UC Urban Bee Lab taxonomist, Amanda Niemela, independently confirmed all of these identifications. All photographic vouchers of specimens were uploaded to an iNaturalist project [https://www.inaturalist.org/projects/ucb-oxford-tract-native-bees].

### Bipartite Network

Plant–pollinator network data were generated using field measurements of floral abundance, non-lethal bee collections, and standardized observational surveys. All interaction data were compiled into a species-by-species interaction matrix, with each plant and pollinator species represented and the observed visitation frequency recorded. Data cleaning and visualization were conducted in RStudio (R version 4.4.0) (R Core Team, 2022) using the tidyverse (Wickham et al., 2019), dplyr (Wickham & Francois, 2015), tidyr (Wickham & Henry, 2020), and ggplot2 (Wickham, 2016) packages. An interaction matrix was generated by amalgamating surveys across days and time points, with each node representing a species and each link representing an observed interaction (Bascompte & Jordano, 2007). Each value in the matrix indicates the frequency of interactions as the number of times that taxa was collected from that flower (Vázquez et al., 2009). This matrix was used to construct quantitative bipartite networks and to compute network and species-level metrics using the bipartite package (Dormann et al., 2017).

To test differences in the plant pollinator network through time, we generated three networks for each time block (morning, midday, afternoon). A fourth matrix, combining all interaction events, was also constructed to characterize the overall network. Species-level visitation strength was calculated for each group as the total number of visits per plant species by summing interaction frequencies for each taxa. Standard network-level indices were calculated for the temporal and overall network, including connectance (Dunne et al., 2002), nestedness metric based on overlap and decreasing fill (NODF) (Almeida-Neto et al., 2008), linkage density (Bersier et al., 2002), H2′ specialization (Blüthgen et al., 2006), modularity (Dormann & Straub, 2014) and evenness (Jost, 2006) using the bipartite package (Dormann et al., 2017). We used permutation tests to generate null models to quantify the differences between all the garden’s network metrics in comparison to the values expected if the networks were random. This was done with swap randomization holding constant sampling intensity (Blüthgen et al., 2006). For each time block, 1,000 null matrices were generated and the network metrics were recalculated to produce a null distribution. Connectance, however, was not tested against the null due to the limitations of the swap algorithm. Observed values were summarized using the null mean, 95% confidence interval, and standard effect sizes (SES). Statistical significance was reported relative to permuted matrices as two-sided p-values. We assessed whether plant traits predicted visitation by honey bees in comparison to native bees using linear regression, with visitation frequency as the response variable and plant traits (inflorescence abundance, corolla width, native status) as predictors, and used statistical correction for multiple hypothesis testing.

## Results

### Network Level Metrics

In total, we directly observed one non-native pollinator (*A. mellifera*) and nine native bees identified to the lowest possible taxonomic order (*B. vosnesenskii, Xylocopa* sp., *Melissodes* sp. (Latreille, 1829), *Anthophora urbana* (Cresson, 1878), *Megachile spp*., *Coelioxys rufitarsis* (Smith, 1854), *Agapostemon subtilior* (Portman, 2024), *Lasioglossum* spp. (Curtis, 1833), and *Halictus tripartitus* (Cockerell, 1895). However, two apparent species of *Megachile* were present in the garden, and one of them is likely the European alfalfa leafcutter bee (*Megachile rotundata*), introduced to the US in the 1940s (Pitts-Singer & Cane 2011). From our total observations (Table S1), we generated a bipartite plant-pollinator network of a single native pollinator garden in the summer of 2025 (Fig. 2).

**Figure 2.**
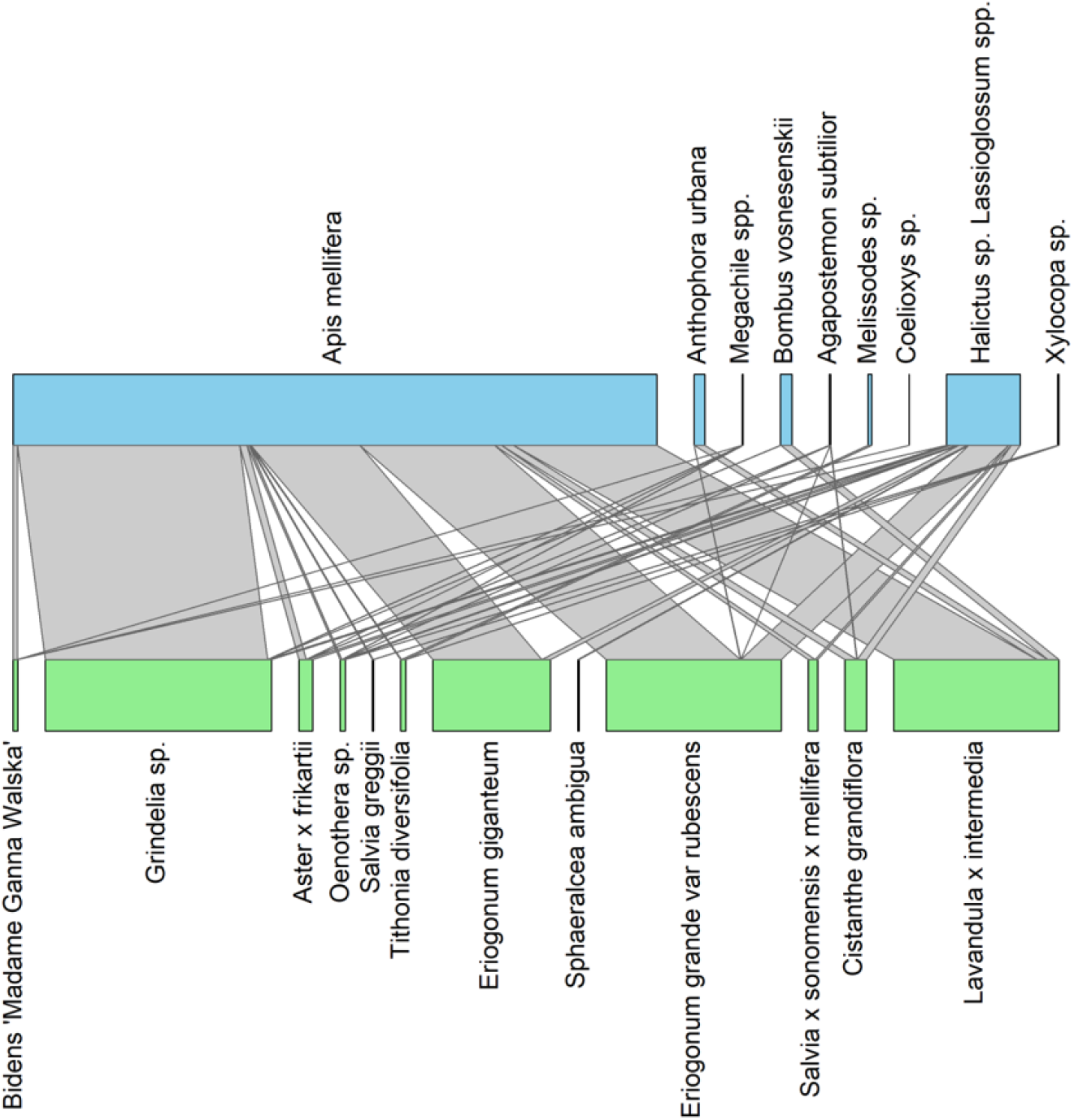
Bipartite plant–pollinator interaction network. The observed interactions between plant species (green, bottom) and bee genera/species (blue, top) within the garden community. The width of each bar represents the total number of interactions, and the lines connecting groups indicate the frequency of interactions between each plant–pollinator pair.

To assess the network structure and quantify the patterns of ecological interactions, we calculated a series of network metrics across the three time points sampled (morning, midday, afternoon) and for the overall network. Nestedness (NODF), specialization (H2’), linkage density, interaction evenness, and modularity are all reported in the supplement (Table S2). Standard effect sizes (SES) were calculated for each metric as deviations from the null expectations for the three temporal networks and the overall network (Fig. 3). Throughout the observation period, the network exhibited a moderate level of connectance (0.34), defined as the fraction of realized interactions relative to the total possible interactions (Dunne et al., 2002). Interaction nestedness, or the proportion of specialist interactions that are a subset of existing generalist interactions, was significantly lower than expected by chance (NODF = 53.73, SES = -2.1, p = 0.04). The overall network-level of specialization (H2’) found in the network was significantly greater than expected by chance (H’=0.39, SES = 4.1, p = 0.006), indicating more specialized interactions than the null model. Linkage density indicated that the average number of links per species was approximately three interactions and was significantly less than expected under random models (SES = -3.78, p = 0.008). This means that there were fewer interactions per species than expected if the network were random. Network evenness was significantly less than expected by chance (0.44, SES = -4.13, p = 0.006). This low evenness value highlighted that interactions were dominated by a few species rather than being distributed throughout the network. Modularity was measured to determine whether certain groups of species interacted more with one another than with other species. Although the modularity score was modest (Q=0.13), given the species interaction frequencies, it was much more highly modular than expected by chance (SES = 5.03, p < 0.001). In total, the overall network indicates strongly non-random structure and greater compartmentalization than expected by chance, with certain species dominating the network through uneven selective interactions. When the network was partitioned into morning, midday, and afternoon time blocks, most metrics were relatively consistent, indicating that the network structure remained stable over time (Fig. 3). However, nestedness differed throughout the day. The morning network did not differ significantly from null model predictions (NODF = 41.04, SES = 0.09, p = 0.88), whereas, both midday (NODF = 27.45, SES = -3.08, p = 0.004) and afternoon networks (NODF = 30.72, SES = -2.41, p = 0.03) exhibited significantly lower nestedness than expected by chance.

**Figure 3.**
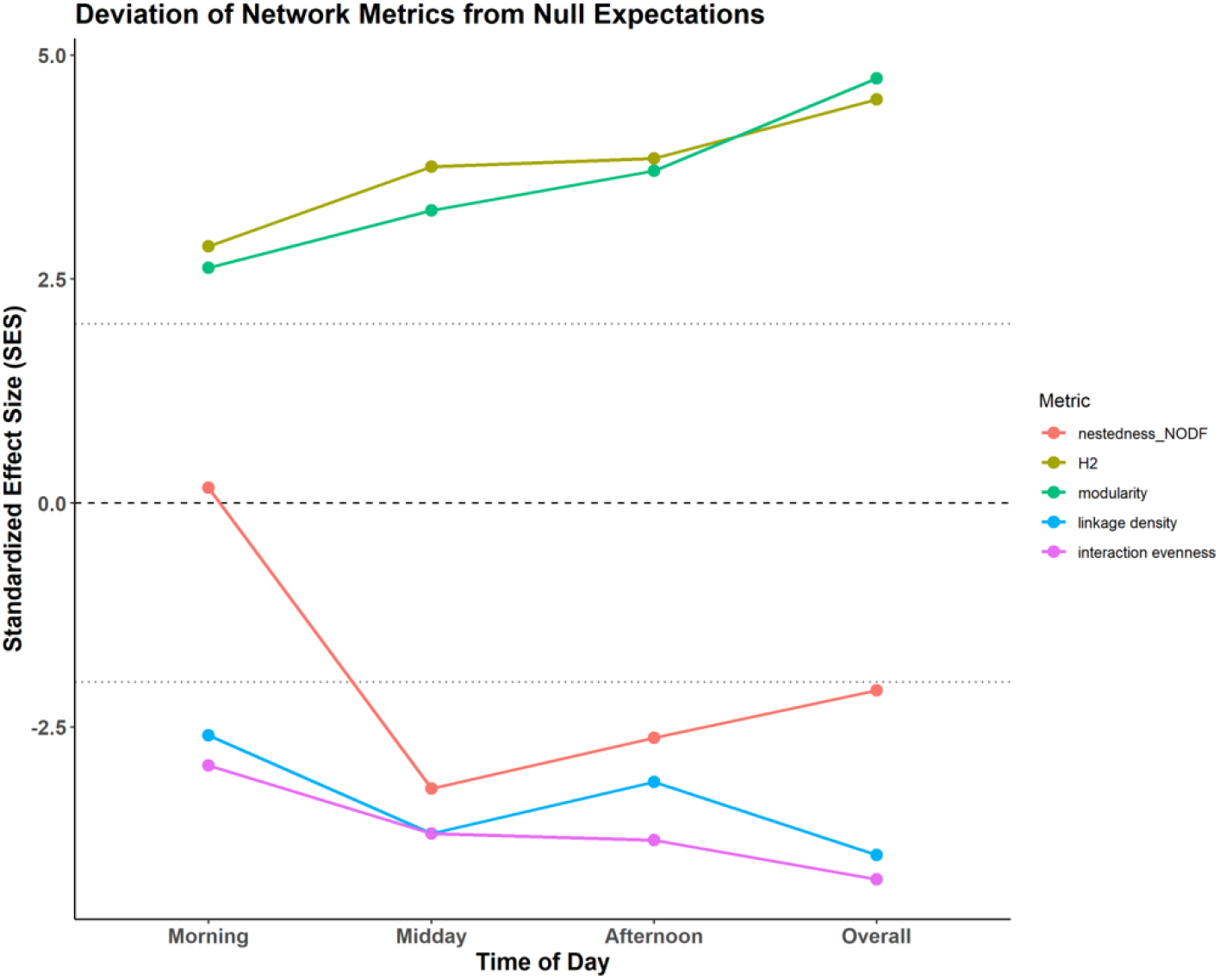
Temporal deviation of network structure relative to null expectations. The standardized effect sizes (SES) of network metrics (nestedness, H_2_ specialization, modularity, linkage density, and interaction evenness) were plotted on a line plot. SES values represent deviation from null model expectations, with the dashed line at 0 representing random and the dotted lines representing significance thresholds.

### Species-Level Metrics

We assessed interaction strength as the frequency of interactions between plant and bee (Fig. 4). The pollinator with the most documented visits was the honey bee *(A. mellifera*). The next most frequently observed pollinators that visited the most flowers were the sweat bees in the family Halictidae (genus: *Halictus & Lasioglossum*), followed by the Apidae family bees of bumblebees (genus: *Bombus)* and the solitary digger bee (*A. urbana)*. We observe, however, that this network is dominated by the non-native honey bee, *A. mellifera*, visiting four main floral species: *Grindelia* spp., *Erigonum grande, Lavandula x intermedia*, and *Eriogonum giganteum*, all of which exhibit the highest visitation frequencies in the network. The other bees that visit a diversity of floral resources, with *Eriogonum* having the highest number of native bee visits.

**Figure 4.**
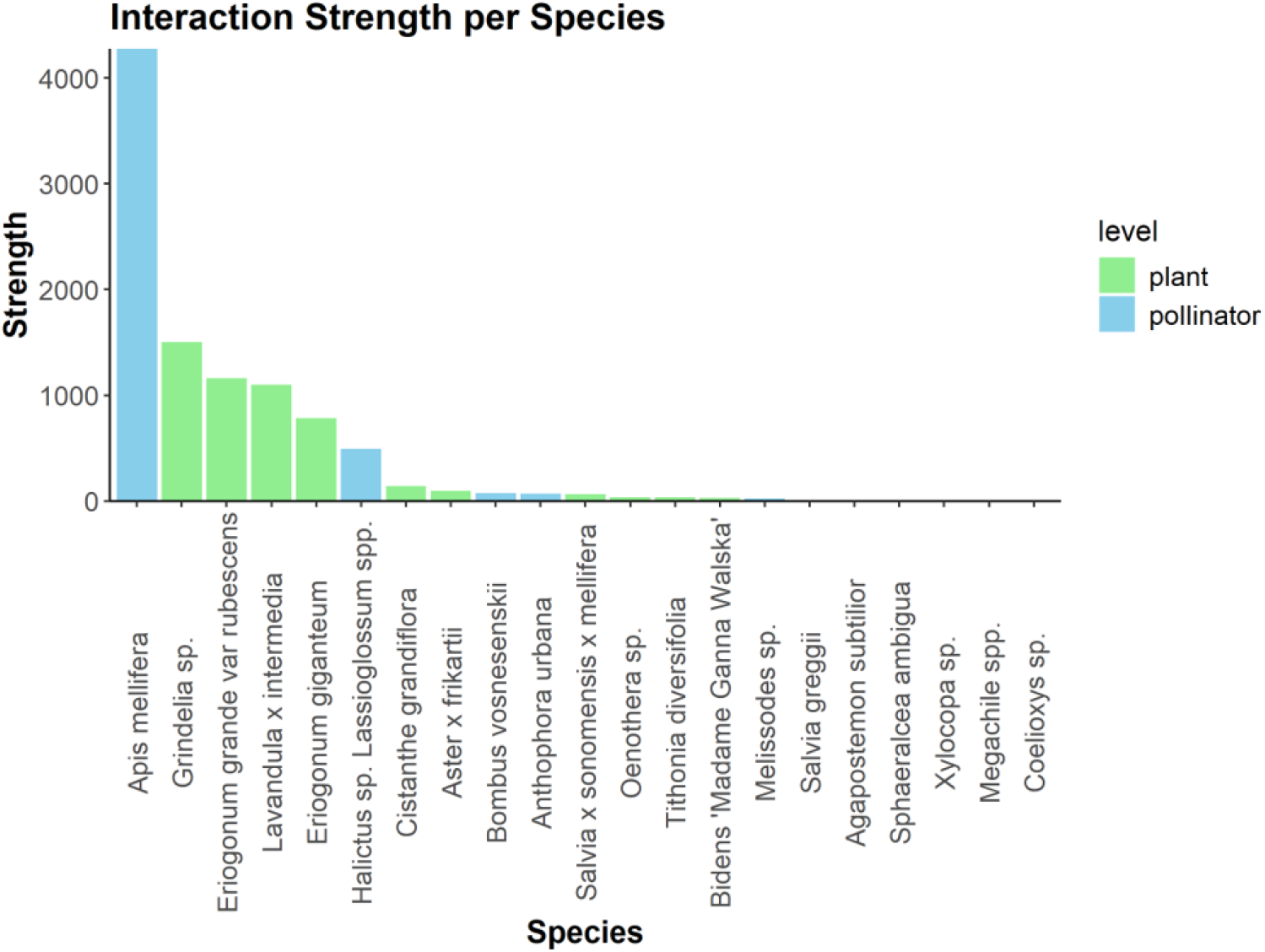
Interaction strength of each focal species in the overall plant-pollinator network across all time points. Each bar depicts the total interaction strength for each species, calculated as the sum of all observed interactions between bee pollinators (blue) and plants (green). This highlights network asymmetry through the dominance of a few highly interacting species and the long tail of taxa with fewer visitations.

Species-level metrics revealed strong asymmetries in visitation patterns (Table S3). *A. mellifera* exhibited the highest degree (11), interaction strength (7.50), and weighted betweenness (1), indicating a dominant and central role within the network. Sweat bees (*Halictus and Lasioglossum*) were the native bees that interacted with the most plants in the network (degree = 9) or 75% of the plants available. Sweat bees also had the highest interaction strength (2.97) of the native bees. We examined species specialization (d’), which quantifies the deviation of a species interaction from random expectations and indicates the degree of generalism versus specialism. *A. mellifera* was a generalist in the network (d’ = 0.1), whereas *Xylocopa* and *Melisodes* were more specialized pollinators (d’ = 0.65 and 0.95, respectively). Most plants showed generalist interaction patterns, except *Sphaeralcea ambigua*, which showed moderate specialization (d’=0.36), and *Tithonia diversifolia*, which showed specialized interactions despite low visitation rates (d’ = 0.78). This was driven by interactions with *Melissodes*, which foraged only on *T. diversifolia*.

To examine patterns of interaction structure, we visualized the network as a heat map, with the color intensity of each cell representing the frequency of interactions between species on a log scale (Fig. 5). Many native bee species exhibited specialized floral interactions, foraging only from a few select flowers. Highly connected plant species included *Grindelia* sp., *Erigonum grande var. rubescens, and Lavandula x intermedia*, with frequent interactions among many pollinator species. *A. mellifera* shows high interaction frequencies, visiting nearly all plant species at a high level. Some plants exhibit specialist interactions, interacting with only one or two pollinators. This includes *Sphaeralcea ambigua*. Similarly, *Melissodes* sp. (Latreille, 1829) and *C. rufitarsis* exhibit specialist interactions, interacting with few plant species. Many bees visit some plants, but not all plants are visited by many bees.

**Figure 5.**
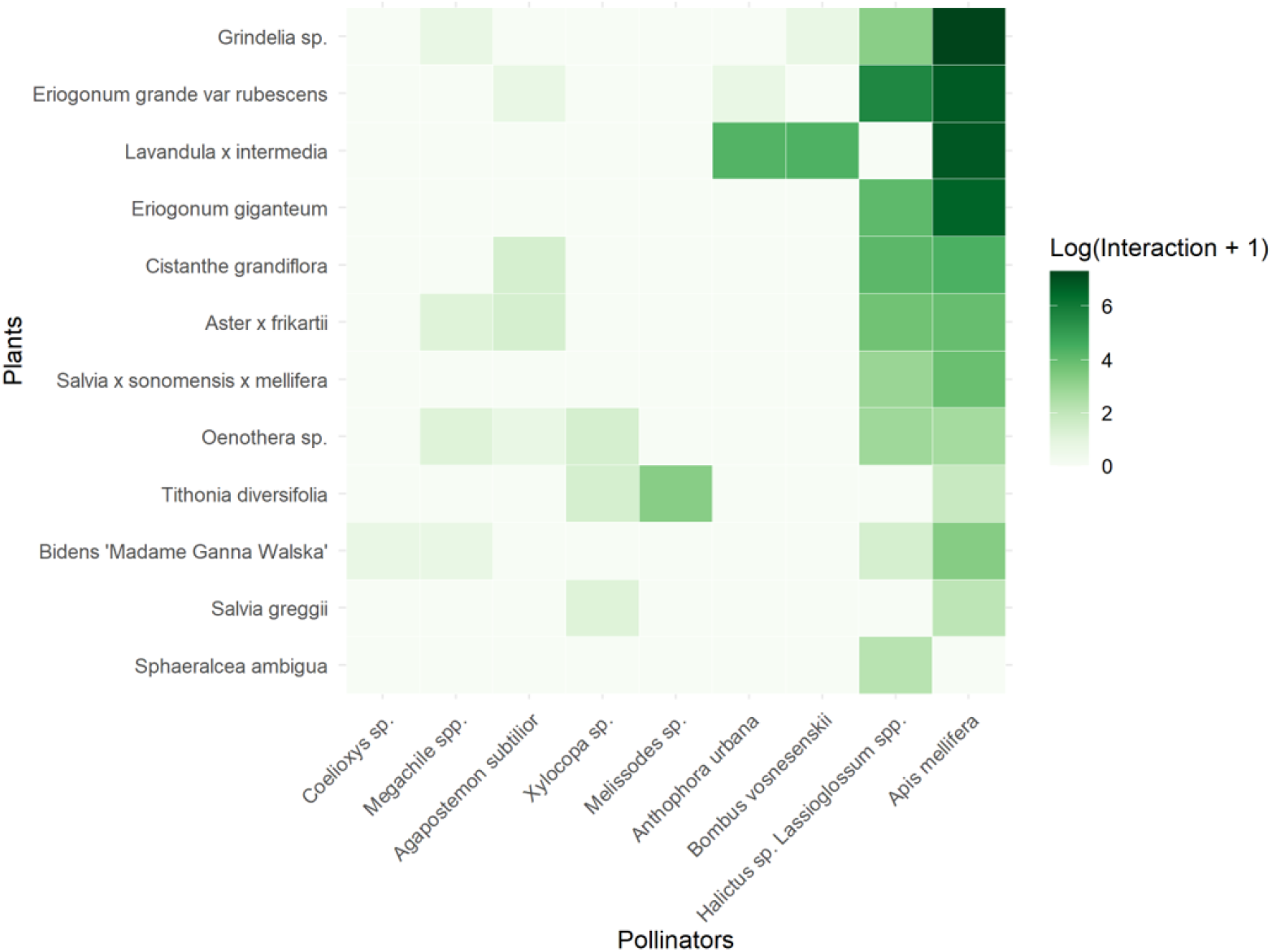
Log-transformed interaction matrix for the pollinator (x-axis) and plant (y-axis) community as a heatmap. The fill color represents the logarithm of interaction strength, with darker shades indicating higher interaction frequencies. White indicates gaps and may indicate that pollinators are not pollinating these plants or that there is a sampling gap.

### Trait-Based Analysis

The relationship between pollinator visitation and floral abundance, or the average number of inflorescence per plant, was assessed, with comparisons between honey bees and native bees (Fig. 6). Honey bee visitation increased significantly with floral abundance (*p* = 0.034), where abundance explained approximately 37% of the variance (β = 1.95 ± 0.80 SE, R^2^ = 0.37). On the other hand, native bee visitation showed no significant relationship to abundance (β = 0.10 ± 0.15 SE, *p* = 0.536, R^2^ = 0.04) (Table S4). *Grindelia* sp. had an average inflorescence count of 359.33 and experienced the highest honeybee visitor count of all plants, with 1479 visitations. Similarly, *Lavandula x intermedia* attracted 953 *A. mellifera* visitors, averaging 497 per inflorescence. *Eriogonum grande var. rubescens* had 894 *A. mellifera* visitations with an average inflorescence count of 96. No other traits showed significant differences between honey and native bees.

**Figure 6.**
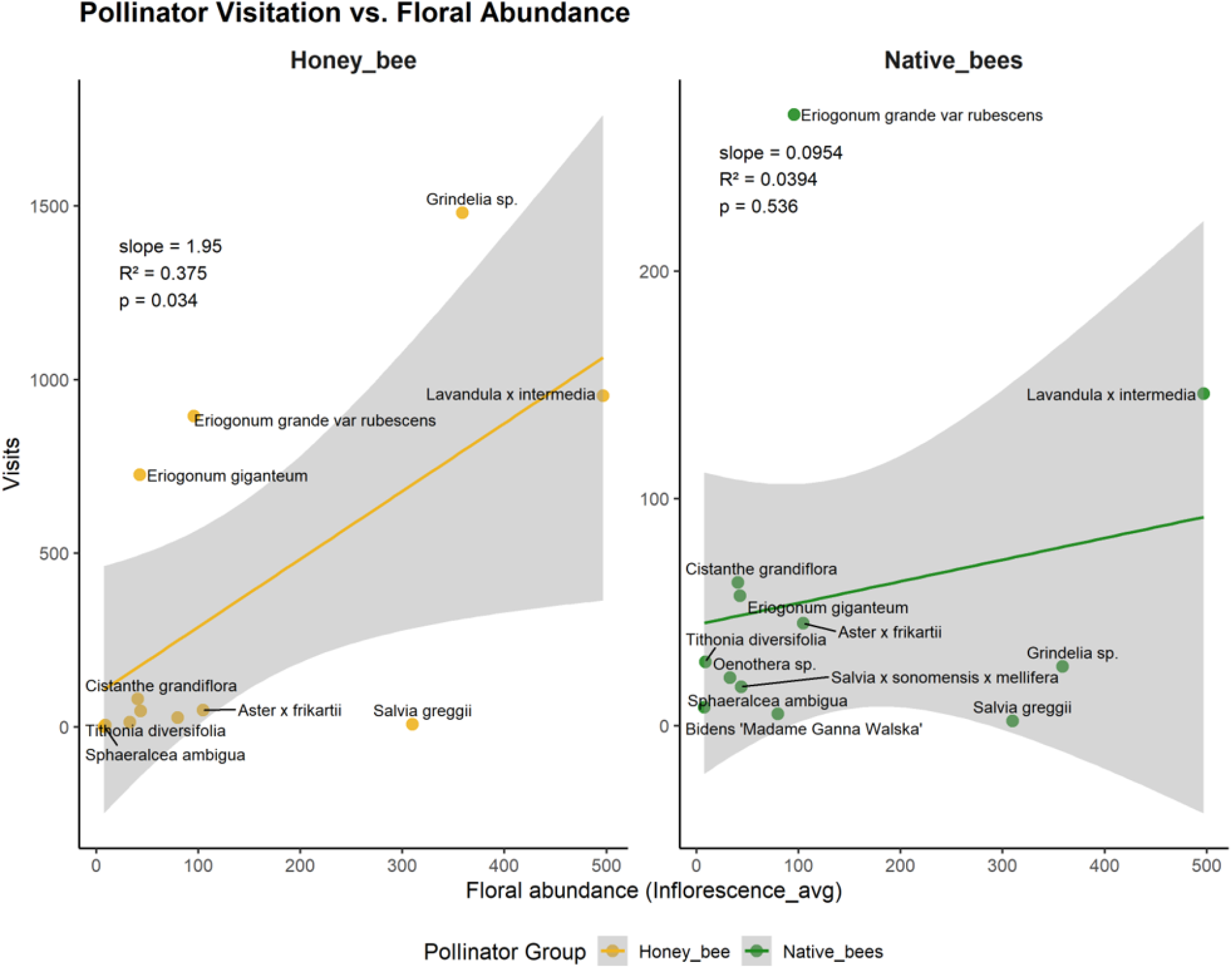
Pollinator visitation as a function of floral abundance for honey bees (yellow, left) and native bees (green, right). Scatterplots show the relationship between floral abundance (average number of inflorescences per plant species) and visitation frequency. Linear regressions were fitted separately for honey bees and native bees. Points are annotated with plant species names. Confidence intervals are shown as grey shadows around each group-specific regression. The honey bee model shows a significant relationship between visitation and floral abundance (slope = 1.95, R2 = 0.375, *p* = 0.034).

## Discussion

This study quantified the plant pollinator interactions in a local campus native bee garden in July of 2025 through non-lethal photography based surveys. We report a non-random bipartite plant-pollinator network with significant modularity and a higher level of specialization than expected by chance. The network is unevenly distributed with dominance from *A. mellifera* and *H. tripartitus*. Early in the day, network nestedness was random. Nestedness decreased across the day, indicating less overlap of generalist and specialist interactions than random. As the day progressed, specialist pollinators interacted with plants outside of what were being primarily used by generalists. This is in contrast to findings found in studies comparing plant-bee networks in urban versus natural areas. Urban networks were more nested, thus meaning greater overlap between generalists and specialists and had greater patterns of generalist foraging (Sirohi et al. 2022). Schmack and Egerer (2023) also found that nestedness increased with increases in floral richness in urban gardens.

Honey bees (*A. mellifera*) exhibited high connectivity within the network, visiting every species except *Sphaeralcea ambigua* (Fig. 1). This indicates that they are a key generalist pollinator in the gardens, but it also raises concerns that they may outcompete native species or serve as a hub for disease transmission within the network (Graystock et al., 2016). In addition to high connectivity in the network, *A. mellifera* also exhibited high interaction strength (Fig. 2, 4). Native bee species demonstrated specialized floral interactions, in which they foraged only from a few select flowers (Fig. 5). *Melissodes* sp. exhibits highly specialized interactions with *Tithonia diversifolia*, as the pollinator was not found on any other plant species. Most native bee species, including *Coelioxys* sp., *Megachile* spp., *A. subtilior, Xylocopa* sp., *Melissodes* sp., *A. urbana*, and *Bombus* spp. displayed more evidence of floral specialization, each foraging on up to four plants. In contrast, native Halictid bees (*Halictus* sp. & *Lasioglossum* sp.) were generalists with foraging preferences most similar to *A. mellifera* (Fig. 5). However, Halictid bees had lower network interaction strength than *A. mellifera* (Fig. 4). Martins et al. (2017) found that bees in urban gardens could be both more specialized and more generalist than bees in semi-natural habitats due to the floral availability, including that of non-native plants.

Modularity between trait-sharing bees and/or more closely related bees in our study was insignificant. There were no significant findings or patterns in floral measurements or in bee traits, including sociality, mean proboscis length, foraging habits, native status, and phenology. We believe that with increased sampling effort, we would expect significant patterns to emerge between floral traits and visitation by native versus non-native bees. However, there was a positive correlation between inflorescence abundance and honey bee visitation. Honeybees had especially high visitation to *Grindelia* sp. and *Lavandula x intermedia*, and overall fewer visits to flowers with fewer inflorescences. These plants likely function as generalists, providing core resources within the network. This is consistent with previous work, such as that by Hung et al. (2019), which suggests that at high floral abundance, honey bee visitation rates increase at a greater rate than those of non-honey bees. There was no significant relationship between native bee visitation and floral abundance (Fig. 6).

Since the urban garden has been curated to best support native bees, we assumed that more than 50% of pollination events would involve native bee species. However, we found that *A. mellifera* accounted for approximately 86% of all floral visitations recorded in this study (Table S1). Non-honey bee interactions were approximately 14% of all floral visitations in the study. Excluding the introduced *Megachile rotundata*, native bee visitation is less than 13.84% of all recorded pollination events, meaning that native bees accounted for fewer than one in seven floral visits.While native species generally exhibited specialist tendencies in our system, the native sweat bee *H. tripartitus* was an exception to this pattern and was a generalist forager. Native bees like *Bombus spp*., *A. urbana*, and *Melissodes sp*. interacted with only a few flower species available during the survey. On the other hand, *A. mellifera*, a non-native pollinator, was recorded visiting all flower species except *Sphaeralcea ambigua. A. mellifera* often makes multiple visits to flowers of the same species before moving on to another flower type (Hung et al. 2019). With a positive correlation between honeybee populations and inflorescence, abundant flowers may have attracted honeybee hives hoping to forage efficiently. Perhaps *A. mellifera* eusociality enables communication with large, accessible masses of the same flower type present at the field site. Hung et al. (2019) observed a positive correlation between *A. mellifera* visitation and floral abundance above a ‘higher flower abundance’ threshold. Garden stewardship for native bees should avoid high abundance of any flower beyond a certain threshold to prevent an excess of honey bees and to avoid direct competition for floral resources. Increasing floral richness through a higher diversity of flowering species may help attract a greater diversity of native bees (Baldock et al., 2019), including flowers with smaller mean sizes (Philpott et al. 2025). Additionally, increasing the size of gardens has been shown to increase abundance and diversity of bee pollinators along the California central coast (Quistberg et al. 2016). Future studies may aim to better understand ‘higher flower abundance’ thresholds for any native bee garden space. Replicating the study’s methods at additional field sites and on additional sampling days may yield more robust and conclusive data on bee traits and floral measurements. Floral abundance data from multiple field sites can clarify the role of ‘higher flower abundance’ thresholds in urban pollinator gardens. This must be done carefully as other studies based in Europe have found that floral abundance does not impact pollinator visitation networks (Schmack & Egerer, 2023) and studies in central California have found gardens with higher floral abundance had greater species richness and abundance of bees (Quitsberg et al., 2016).

We compared our findings to that of the past surveys of the research farms previous pollinator garden. All full-season bees reported in the 2004 study, including *A. mellifera, A. subtilior, A. urbana, Megachile perihirta* (Cockerell 1898), *M. rotundata*, and *Halictus tripartitus*, were found in our study, suggesting that the phenology and range of the urbanized bees remain since the early 2000s. In fact, the garden provided habitat for a variety of life history strategies, including social, solitary, and parasitic bees. *A. mellifera* and *B. vosnesenskii* are social bees with division of labor in their nests (Frankie et al. 2009). Other bees found in our study are solitary, such as *A. urbana, M. perihirta, M. rotundata, A. subtilior, Melissodes* sp., *C. rufitarsis, H. tripartitus, Lasioglossum* sp., and *Xylocopa* sp. This group of bees includes both ground and cavity nesters, suggesting that the garden is providing the necessary nesting habitat for these species. This may be partially attributed to the garden’s bare soil and the lack of mulch and grasses to allow for burrowing opportunities. The garden also supported a parasitic bee (Frankie et al. 2009). *Megachile spp*. acts as the host for the parasitic bee, *C. rufitarsis. Coelioxys* is a genus of cleptoparasitic bees, which do not possess the characteristic hairs of most bees and instead appear more wasp-like. Female *Coelioxys* do not collect pollen or nectar provisions for their offspring, instead this cleptoparasite will lay eggs in a host bee species’ nest so that the parasitic offspring can hatch and eat the pollen provisions provided by the host species to their offspring. As this parasitic interaction depends on a sufficient host population, it may serve as an indicator of the health of the *Megachile* population at the study site (Scott et al. 2000). Both host and parasite were identified in our study and in a 2004 study of the original pollinator garden in Berkeley, including records of *C. rufitarsis* in July 2004 (Wojcik et al. 2008).

We report two species of the genus *Megachile:* the non-native, alfalfa leafcutter, *Megachile rotundata*, and the native, Western leafcutter, *Megachile perihirta. M. rotundata* could be found widely in the US in the 1940s and was introduced to the Americas from Eurasia to pollinate alfalfa. While most *Megachile* species are efficient pollinators of alfalfa, native *Megachile* were not found in sufficient numbers to meet the increasing demand for alfalfa as cattle feed (Pitts-Singer & Cane 2011). Today, *Megachile rotundata* and *Megachile perihirta* are urbanized, full-season bees (Fig. 1). In the 2004 season at the Oxford Tract, 29 individuals of *M. rotundata* and 70 individuals of *M. perihirta* were collected with pan traps. On one sampling day in July of 2004, 17 *M. perihirta* and two *M. rotundata* individuals were collected by a pan trap (Wojcik et al. 2008). Across three sampling days in July 2025, there were six pollination events by *Megachile* spp. In addition to *M. perihirta* and *M. rotundata*, the 2004 study reported three *individuals of M. brevis onobrychidis* (Cresson 1908) in pan traps in July (Wojcik et al. 2008). Discrepancies may be attributed to a decrease in garden size between the original and new Oxford Tract bee gardens. Additionally, the loss of *Megachile* diversity may be attributed to pathogen spillover from both non-native and native Megachile species via shared floral resources. Studies of Megachilidae (genus *Osmia* (Panzer, 1806)) have shown that non-native megachilid species can introduce new diseases, such as chalk-brood fungus, to the nests of native megachilid populations (LeCroy et al. 2022).

Woijick et al. (2008) reported one *Bombus vosnesenskii* pan trap collection in July of 2004, as *Bombus* peaks in spring and declines by fall. Our study reports 77 pollination events by *Bombus spp*. While bumblebees are considered generalists and perform well in urban environments, their low abundance in this study aligns with previous studies of the Oxford Tract garden and known *Bombus* phenology (McFrederick & LeBuhn 2006; Wojcik et al. 2008). In contrast to previous studies, *Melissodes* sp. exhibits extreme specialist behavior, foraging on only one plant species, *Tithonia diversifolia. Melissodes* spp. have been reported to forage on another sunflower species (*Helianthus annuus*), Sulphur Cosmos (*Cosmos sulphureus)*, Mexican aster (*Cosmos bipinnatus*), and black-eyed Susan (*Rudbeckia hirta)* (Frankie et al. 2009). In fact, *Melissodes* sp. was described as the primary visitor of golden tickseed (*Coreopsis tinctoria)* (Frankie et al. 2005).

*Melissodes lupina* (Cresson 1878) and *Melissodes robustior* (Cockerell 1915), species previously found in the Oxford Tract garden, are described as specialists to the entire daisy family, Asteraceae (Frankie et al. 2005; Wojcik et al. 2008). It is likely that our field site provided limited floral resources for Melissodes, resulting in a conditional specialist behavior. *Melissodes* are late spring-early summer (Frankie et al. 2005). In July of 2004, the Oxford Tract garden recorded 16 individuals of Melissodes robustior in pan traps (Wojcik et al. 2008). Our study reports 25 pollination events by *Melissodes* sp. In fact, a *Melissodes* sp. male was observed to be engaged in a territorial flight pattern, hovering from bloom to bloom on *Tithonia diversifolia*, with occasional copulation attempts with a female *Melissodes* sp.

The most abundant native bees in our study were the sweat bees (Family: Halictidae), *H. tripartitus* and *Lasioglossum sp*. Both of which exhibited generalist foraging behaviors (Fig. 1). *H. tripartitus* and *Lasioglossum sp*. were found on all but three plant species: *Lavandula x intermedia, Salvia greggii*, and *Tithonia diversifolia*. We observed 491 pollination events by *H. tripartitus* and *Lasioglossum sp*. In the 2004 study at the Oxford Tract, the July pan-trap collections contained 15 *H. tripartitus*, six *Lasioglossum incompletum* (Crawford, 1907), and one *Halictus rubicundus* (Christ, 1791). As Halictidae populations peak in late spring to early summer, their abundance at the time of this study is unknown and requires more observations over time (Wojcik et al. 2008). Large populations of halictids confirm the use of bare soil with small amounts of natural leaf litter in gardens to provide habitat for ground-nesting bees. We also observed eight pollination events by *A. subtilior* (Fig. 1). In July 2004, there were no *A. subtilior* individuals in the pan trap collection. *A. subtilior* is a full-season bee with multi-generational emergences throughout the year. In other words, *A. subtilior* will emerge from their nesting cavities multiple times throughout the season (Frankie et al. 2019). It is likely that the 2004 study was conducted at a point between emergences of new individuals from their nest cavities, resulting in a lower number of individuals. Alternatively, there may have been a shift in the phenology of *A. subtilior*, with an earlier emergence. We observe that bees prefer to forage at specific times of day, as suggested by Frankie et al. (2009). Species diversity was higher during the midday and afternoon periods than in the morning. However, *A. mellifera* showed increased pollination events in the morning and midday.

Similarly to the 2009 study of urban California gardens, *Salvia greggii* in our study attracted a few large carpenter bees *(Xylocopa* sp.; 2 visits) and honey bees *(A. mellifera)* (Frankie et al. 2009). Honey bees had lower visitation rates on *Salvia greggii* than those of other plant species, with only seven visits in our study. *Lavandula × intermedia* attracted *A. mellifera* and *A. urbana*, as described by Frankie et al. (2009). There were no visitations by *Xylocopa* sp. or Megachilidae bees on *Lavandula x intermedia*, which were previously described to have low visitation rates. Species present in the garden but not observed during the formal survey times included *Anthidium* sp. and *Bombus* sp. 2. Studies of the Oxford Tract Garden from July 2004 reported one *Anthidium maculosum* (Cresson 1878) in pan traps. Finally, we do not report any occurrences of pollination events by the cavity-nesting genus *Hylaeus* (Frankie et al. 2005). One *Hylaeus rudbeckiae* (Cockerell & Casad, 1895) individual was reported in July 2004 from a pan trap (Woijcik 2008). As *Hylaeus* is active from June to October, with peak abundance in September, a low presence of *Hylaeus* was expected (Woijcik 2008).

Conversion of natural habitat to urban areas may significantly limit native bee populations (Hernandez et al. 2009). Fortunately, urban gardens have been shown to be successful in supporting the diversity of bees amidst environmental threats (McFrederick and LeBuhn 2006; Philpott et al., 2025). With proper techniques and practices like reconciliation ecology, we may find solutions to return urban lands to bees and support their diversity of forms (McFrederick and LeBuhn 2006). While pollinators in urban gardens may face concerns around competition for floral resources from managed honey bees (Casanellas abella et al., 2025), we may be able to decrease the chance of negative impacts through providing abundance. This can be through increasing plant diversity, including plants that flower at different times of the year, and avoiding dominance of any one plant species. Valido et al. (2019) found when a single non-native generalist dominates a network, it alters overall pollinator functionality and can inadvertently depress the broader ecosystem services required to maintain wildland and urban plant diversity. When we design native pollinator gardens, we must consider the diversity of life history strategies, foraging behaviors, and phenological timing of emergence expressed by native bees and their connections with floral preferences, in order to support the highest species richness possible. Local factors will likely be critical drivers of pollinator populations, therefore gardeners have the ability to promote native bee conservation through informed management practices (Quistberg et al., 2016). Increasing the size of gardens, planting a diversity of different flower species that bloom throughout different seasons of the year, providing bare ground for nesting resources and using mulch sparingly are all helpful avenues of providing refuge for native bees amongst the impacts of urbanization.

## Acknowledgments

IN thanks the Rose Hills Foundation and the Berkeley Food Institute for financial support. The authors would like to thank the University of California, Berkeley, the Urban Bee Lab, and Dr. Gordon W. Frankie for providing access to the field site. Special thanks to Amanda Niemela, taxonomist of the UC Berkeley Urban Bee Lab, for bee taxonomy and identification support and to Emma Coflin for plant identification support.

