## Supplemental Tables for "Pollinator Plant Network Interactions of Bees (Hymenoptera: Anthophila) in a California Urban Garden"

Supplementary Table 1: A matrix of the overall plant–pollinator interactions across the morning, midday, and afternoon. Each value indicates a visitation of a bee taxa (columns) on the flowering parts of the plant varietals (rows).

| **Bees:** | **Apidae** | **Apidae** | **Apidae** | **Apidae** | **Megachilidae** | **Megachilidae** | **Halictidae** | **Halictidae** |
|---|---|---|---|---|---|---|---|---|
| **Plants:** | *Apis mellifera* | *Anthophora urbana* | *Bombus vosnesenskii* | *Melissodes robustior* | *Megachile sp.* | *Coelioxys rufitarsis* | *Agapostemon subtilior* | *H. tripartitus & Lasioglossum sp.* |
| ***Bidens 'Madame Ganna Walska'*** | 26 | 0 | 0 | 0 | 1 | 1 | 0 | 3 |
| ***Grindelia sp.*** | 1479 | 0 | 1 | 0 | 1 | 0 | 0 | 24 |
| ***Aster x frikartii*** | 48 | 0 | 0 | 0 | 2 | 0 | 3 | 40 |
| ***Oenothera sp.*** | 13 | 0 | 0 | 0 | 2 | 0 | 1 | 15 |
| ***Salvia greggii*** | 7 | 0 | 0 | 0 | 0 | 0 | 0 | 0 |
| ***Tithonia diversifolia*** | 5 | 0 | 0 | 25 | 0 | 0 | 0 | 0 |
| ***Eriogonum giganteum*** | 725 | 0 | 0 | 0 | 0 | 0 | 0 | 57 |
| ***Sphaeralcea ambigua*** | 0 | 0 | 0 | 0 | 0 | 0 | 0 | 8 |
| ***Eriogonum grande var rubescens*** | 894 | 1 | 0 | 0 | 0 | 0 | 1 | 267 |
| ***Salvia x sonomensis x mellifera*** | 46 | 0 | 0 | 0 | 0 | 0 | 0 | 17 |
| ***Cistanthe grandiflora*** | 80 | 0 | 0 | 0 | 0 | 0 | 3 | 60 |
| ***Lavandula x intermedia*** | 953 | 70 | 76 | 0 | 0 | 0 | 0 | 0 |
| ***Total visits*** | 4276 | 71 | 77 | 25 | 6 | 1 | 8 | 491 |
| ***Non-honeybee total*** | 679 | | | | | | | |

Supplementary Table 2: Network metrics for the three temporal plant-pollinator networks measured in the morning, midday, and afternoon, as well as the overall network metrics when each of the temporal networks was combined.

| Network | Metric | Observed | Null_mean | Null_ci_low | Null_ci_high | SES | p_two_sided | n_perms |
|---|---|---|---|---|---|---|---|---|
| Morning | nestedness_NODF | 41.036 | 40.687 | 32.213 | 45.098 | 0.097 | 0.882 | 1000.000 |
| Morning | H2 | 0.537 | 0.256 | 0.107 | 0.478 | 2.826 | 0.018 | 1000.000 |
| Morning | linkage density | 3.008 | 3.237 | 3.029 | 3.363 | -2.638 | 0.036 | 1000.000 |
| Morning | evenness | 0.465 | 0.484 | 0.469 | 0.493 | -2.844 | 0.024 | 1000.000 |
| Morning | modularity | 0.092 | 0.052 | 0.028 | 0.087 | 2.574 | 0.020 | 1000.000 |
| Midday | nestedness_NODF | 27.451 | 43.694 | 31.699 | 52.288 | -3.081 | 0.004 | 1000.000 |
| Midday | H2 | 0.483 | 0.179 | 0.077 | 0.368 | 3.752 | 0.006 | 1000.000 |
| Midday | linkage density | 2.777 | 3.179 | 2.912 | 3.330 | -3.481 | 0.010 | 1000.000 |
| Midday | interaction evenness | 0.448 | 0.480 | 0.457 | 0.491 | -3.448 | 0.016 | 1000.000 |
| Midday | modularity | 0.172 | 0.071 | 0.036 | 0.147 | 3.182 | 0.028 | 1000.000 |
| Afternoon | nestedness_NODF | 30.719 | 43.959 | 31.788 | 53.555 | -2.414 | 0.030 | 1000.000 |
| Afternoon | H2 | 0.416 | 0.202 | 0.111 | 0.334 | 3.706 | 0.004 | 1000.000 |
| Afternoon | linkage density | 2.816 | 3.125 | 2.908 | 3.286 | -3.298 | 0.010 | 1000.000 |
| Afternoon | evenness | 0.464 | 0.492 | 0.476 | 0.504 | -3.757 | 0.008 | 1000.000 |

| | | | | | | | | |
|---|---|---|---|---|---|---|---|---|
| Afternoon | modularity | 0.184 | 0.089 | 0.051 | 0.154 | 3.484 | 0.006 | 1000.000 |
| Overall | nestedness_NODF | 53.731 | 66.571 | 54.405 | 77.648 | -2.129 | 0.040 | 1000.000 |
| Overall | H2 | 0.397 | 0.140 | 0.063 | 0.298 | 4.122 | 0.006 | 1000.000 |
| Overall | linkage density | 3.066 | 3.372 | 3.183 | 3.470 | -3.784 | 0.008 | 1000.000 |
| Overall | interaction | 0.438 | 0.461 | 0.447 | 0.468 | -4.129 | 0.006 | 1000.000 |
| Overall | modularity | 0.130 | 0.042 | 0.022 | 0.087 | 5.026 | 0.0001 | 1000.000 |

Supplementary Table 3: Species-level network metrics for the bees (pollinators) and the plants.

| Group | Species | degree | normalised.degree | Species. strength | d' | betweenness | weighted.between | effective.partners |
|---|---|---|---|---|---|---|---|---|
| **Plants** | ***Bidens 'Madame Ganna Walska'*** | 4 | 0.444 | 1.179 | 0.043 | 0.125 | 0 | 1.813 |
| **Plants** | ***Grindelia sp.*** | 4 | 0.444 | 0.574 | 0.081 | 0.125 | 0.642 | 1.097 |
| **Plants** | ***Aster x frikartii*** | 4 | 0.444 | 0.801 | 0.13 | 0.125 | 0 | 2.454 |
| **Plants** | ***Oenothera sp.*** | 5 | 0.556 | 0.867 | 0.2 | 0.125 | 0.17 | 3.364 |
| **Plants** | ***Salvia greggii*** | 2 | 0.222 | 0.252 | 0.155 | 0 | 0 | 1.698 |
| **Plants** | ***Tithonia diversifolia*** | 3 | 0.333 | 1.376 | 0.777 | 0 | 0 | 2.03 |
| **Plants** | ***Eriogonum giganteum*** | 2 | 0.222 | 0.286 | 0.025 | 0.125 | 0 | 1.298 |
| **Plants** | ***Sphaeralcea ambigua*** | 1 | 0.111 | 0.016 | 0.355 | 0 | 0 | 1 |
| **Plants** | ***Eriogonum grande var rubescens*** | 4 | 0.444 | 0.892 | 0.071 | 0.125 | 0.189 | 1.737 |
| **Plants** | ***Salvia x sonomensis x mellifera*** | 2 | 0.222 | 0.045 | 0.032 | 0.125 | 0 | 1.792 |

| | | | | | | | | |
|---|---|---|---|---|---|---|---|---|
| **Plants** | ***Cistanthe grandiflora*** | 3 | 0.333 | 0.5156 | 0.117 | 0.125 | 0 | 2.161 |
| **Plants** | ***Lavandula x intermedia*** | 3 | 0.333 | 2.196 | 0.135 | 0 | 0 | 1.622 |
| **Bees** | ***Apis mellifera*** | 11 | 0.917 | 7.502 | 0.101 | 0.512 | 1 | 4.809 |
| **Bees** | ***Anthophora urbana*** | 2 | 0.167 | 0.065 | 0.335 | 0.018 | 0 | 1.077 |
| **Bees** | ***Megachile spp.*** | 4 | 0.333 | 0.113 | 0.392 | 0.113 | 0 | 3.78 |
| **Bees** | ***Bombus spp.*** | 2 | 0.1667 | 0.07 | 0.34 | 0.017 | 0 | 1.071 |
| **Bees** | ***Agapostemon subtilior*** | 4 | 0.333 | 0.084 | 0.359 | 0.042 | 0 | 3.51 |
| **Bees** | ***Melissodes sp.*** | 1 | 0.083 | 0.758 | 0.947 | 0 | 0 | 1 |
| **Bees** | ***Coelioxys sp.*** | 1 | 0.083 | 0.032 | 0.531 | 0 | 0 | 1 |
| **Bees** | ***Halictus sp. Lasioglossum spp.*** | 9 | 0.75 | 2.976 | 0.323 | 0.19 | 0 | 4.532 |
| **Bees** | ***Xylocopa sp.*** | 3 | 0.25 | 0.4014 | 0.655 | 0.107 | 0 | 2.951 |

Supplementary Table 4: Linear model outputs to test the impact of floral abundance on the visitation of honey bees in comparison to native bees.

| Model Term | Honey Bees | Native Bees |
|---|---|---|
| **Intercept (Estimate)** | 91.64 | 44.33 |
| **Intercept Std. Error** | 163.84 | 30.56 |
| **Intercept t-value** | 0.559 | 1.45 |
| **Intercept p-value** | 0.588 | 0.178 |
| **Slope: Abundance (Estimate)** | 1.955 | 0.095 |
| **Slope Std. Error** | 0.798 | 0.149 |
| **Slope t-value** | 2.448 | 0.641 |
| **Slope p-value** | **0.034*** | 0.536 |
| **Residual Std. Error** | 426.4 | 79.55 |
| **$R^2$** | 0.375 | 0.039 |
| **Adjusted $R^2$** | 0.312 | –0.057 |
| **F-statistic** | 5.994 | 0.411 |
| **F-statistic p-value** | 0.034 | 0.536 |
| **N (plants)** | 12 | 12 |
